# Acute Effects of Cigarette on Endothelial Nitric Oxide Synthase, Vascular Cell Adhesion Molecule 1 and Aortic Intima Media Thickness “Cigarette smoke–induced pro-atherogenic changes”

**DOI:** 10.1101/2021.01.17.426972

**Authors:** Meity Ardiana, Anwar Santoso, Hanestya O. Hermawan, Ricardo A. Nugraha, Budi S. Pikir, I Gde Rurus Suryawan

## Abstract

**Background:** Cigarette smoking could induce endothelial dysfunction and increase of circulating markers of inflammation by activation of monocytes. This can lead to the increased of intima media thickness (IMT) of entire blood vessel and result acceleration of atherosclerosis process. However, to our knowledge, little is known about the role of cigarette smoking in this atherosclerotic inflammatory process.

**Objective:** The aim of this study is to explore the link between cigarette smoking on endothelial nitric oxide synthase (e-NOS) and vascular cell adhesion molecule 1 (VCAM-1).

**Methods:** An experimental study with post-test only controlled group design was used in this study. We used 18 Wistar rats (*Rattus norvegicus*) randomly subdivided into 2 groups, group K (−) were given no tobacco smoking exposed, whereas group K (+) were exposed to 40 cigarettes smokes daily. After 28 days, samples were analyzed for e-NOS, VCAM-1 and aortic IMT.

**Results:** Our results indicate that tobacco smoke can enhance the expression of VCAM-1 on mouse cardiac vascular endothelial cell, resulting in decreased expression of e-NOS level and increased of aortic IMT. Linear regression model found that eNOS level negatively correlated wiith aortic IMT (r^2^ = 0.584, β = −0.764, *p* < 0.001), whereas VCAM-1 expression did not correlate with aortic IMT (r^2^ = 0.197, *p* = 0.065).

**Conclusion:** Low e-NOS level and high VCAM-1 level observed following after cigarette smoke exposure may increase aortic IMT.

**Clinical significance:** Increasing evidence suggests that cigarette smoke exposure could induce VCAM-1 (enhance pro-atherogenic property),and decreased of e-NOS level (anti-atherogenic depletion). Thus, cigarette smoke may represent a significant risk factor for atherosclerosis by increasing aortic IMT. This evidence is discussed herein.

## INTRODUCTION

Cigarette smoking is the most important modifiable risk factor for developing atherosclerosis including cerebrovascular accident, peripheral arterial disease and coronary heart disease^1^. In a meta-analysis from fifty-five eligible studies (43 cross-sectional, 10 cohort and 2 case-control studies), the odds ratio (ORs) of peripheral arterial disease (PAD) associated with cigarette exposed was 2.71 (95% CI: 2.28-3.21; *p*<0.001)^2^. In a meta-analysis from 75 cohorts (2.4 million participants) that adjusted for cardiovascular risk factors other than coronary heart disease, multiple-adjusted pooled ORs of smoking versus non-smoking was 1·25 (95% CI: 1·12–1·39, *p*<0·0001)^3^.

Even though epidemiologic studies clearly stated negative effect of cigarette smoking for cardiovascular diseases, the underlying mechanisms have yet to be confirmed. The pathogenesis and pathophysiologic mechanisms by which exposure to cigarette smoke could accelerate atherosclerosis cardiovascular disease are complex and challenging, due to more than 5000 different mixture chemicals inside the cigarette smoke itself^4^. Several potential contributing factors to atherogenesis inside the cigarette smoke are (1) polycyclic aromatic hydrocarbons, (2) oxidizing agents, (3) particulate matter, and (4) nicotine^5^.

One of the most important factor contributing for pro-atherogenic is nicotine, which has commonly been studied using cigarette smoke condensates^6^. In addition to its role as the habituating agent in tobacco, nicotine also accelerates atherosclerosis cardiovascular disease. There are several potential mechanisms of the pro-atherogenic effects of nicotine: (1) inducing endothelial dysfunction, (2) modifying lipid profile, (3) increasing inflammatory response, (4) inducing the release of catecholamines, which may increases heart rate and blood pressure, (5) increases platelet aggregability, (6) direct actions on the cellular elements participating in plaque formation, and (7) induces the proliferation and migration of vascular smooth muscle cells into the intima, mediated in part by TGFβ ^7^. These pathomechanisms of nicotine could lead to the increases of intimia media thickness of the entire blood vessel, leading to the greater risk of developing atherosclerosis^8^.

To learn deeper about the pathomechanisms of the diseased endothelium, we need to study all the oxidizing, inflammatory, and thrombotic molecules which are not in equilibrium state. In the model of atherosclerosis cardiovascular diseases, a pathological imbalance between prothrombotic and antithrombotic state, prooxidant and antioxidant state, pro-inflammatory and anti-inflammatory state are observed^9^. Considerable evidence supports the importance of inflammation and hypercoagulability to promote atherogenic state^10^. There is abundant literature concerning the role of biomarkers of pathological imbalance in atherosclerosis.

Cell adhesion molecules are the essential pro-inflammatory and pro-atherogenic proteins that represent a hallmark of endothelial dysfunction and atherosclerosis. P-selectin, vascular cell adhesion molecule (VCAM)-1, intercellular adhesion molecule (ICAM)-1, and PECAM-1 were demonstrated to be involved in the formation of atherosclerosis plaque^11^. Beyond the others cell adhesion molecules, VCAM-1 plays as an important factor in neointima proliferation following nicotine-induced arterial injury, an area of research important for atherosclerosis cardiovascular diseases^12^. In the nicotine-induced arterial injury model, VCAM-1 expression is highly induced in the proliferation and migration of neointimal smooth muscle cells^13^.

Previous studies showed that upregulation of endothelial nitric oxide synthase (e-NOS) expression and activity has its important role in the protection of endothelium^14–16^. e-NOS could stimulate endothelium-dependent relaxation and protect against development VCAM-1-induced endothelial dysfunction^17^. However, to our knowledge, little is known about the role of cigarette smoking in this atherosclerotic inflammatory process. This study aims to explore the link between cigarette smoking on e-NOS and VCAM-1, which results to the development of aortic intima media thickness (IMT) of the experimental animals.

## MATERIAL AND METHODS

### Ethics approval

Animal experimental study were conducted under the approval of the Institutional Animal Care and Use Committee of Universitas Airlangga (UNAIR), Surabaya, Indonesia (animal approval no: 2.KE.184.10.2019) under the name of Meity Ardiana as the Principal Investigator. Study was carried out in strict accordance to internationally-accepted standards of the Guide for the Care and Use of Laboratory Animals of the National Institute of Health.

### Animals

The present study used 18 male Wistar rats (*Rattus novergicus*), eight weeks of age (average body weight 150-200 grams). The rats were housed in microisolator cages and maintained in a constant room temperature ranging from 22°C to 25°C, with a 12-h light/12-h dark cycle, under artificially controlled ventilation, with a relative humidity ranging from 50% to 60%. The rats were fed a standard balanced rodent diet and water were provided ad libitum.

### Experimental design and groups

The present study design was a randomized post-test only controlled group design using quantitative method. We extracted 18 male Wistar rats, randomized and then allocated them into 2 groups. Group 1 were given no exposed to tobacco smoke, whilst group 2 were given 40 cigarrete smokes daily for 28 days as seen in **Figure 1**. Each cigarette smoke contains 39 mg of tar and 2.3 mg of nicotine The enrolled subjects were analyzed for vascular cell adhesion molecule 1 (VCAM-1), endothelial nitric oxide synthase (e-NOS), and aortic intima media thickness (IMT) after 28 days of consecutive experiments.

**Figure 1:**
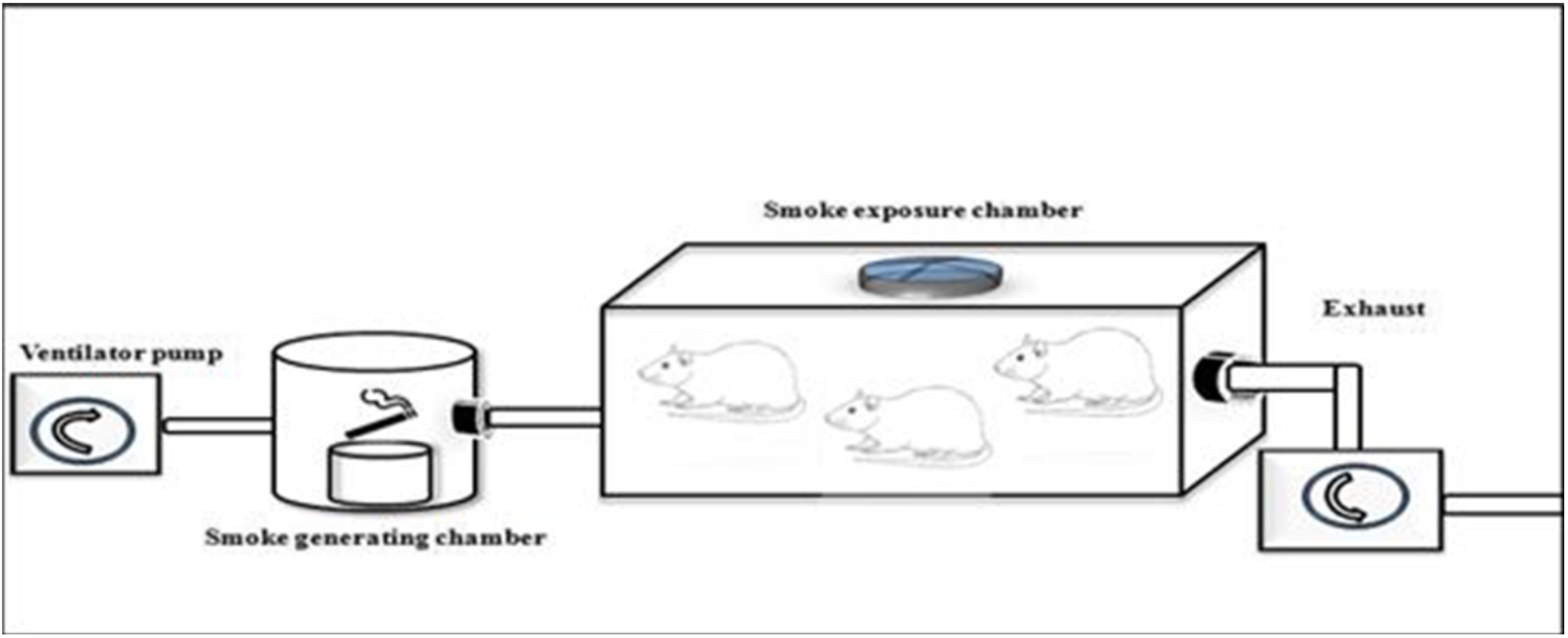
Illustration of how to exposed rats in the K(+) group to the cigarrete smokes. Exposure to tobacco smokes were done using sidestream technique from peristaltic pump, smoke producer chamber, and inhalation chamber, connected by modified silicon tube.

### Aortic Intima Media Thickness (IMT)

Thoracic aortas were prepared as distal aortic arch by cutting from left ventricle. The post mortem samples of descending thoracic aortas obtained by dissection were fixed in 10% formaldehyde, embedded in paraffin, and sectioned at a thickness of 6 μm. The mounted tissues were stained using hematoxylin and eosin. Aortic intima media thickness was measured via Leica DMD 108 (Leica Microsystems GmbH, Wetzlar, Germany). Each sample was measured blindly as “micrometer (μm)” from six different locations of the vessel wall. Arithmetic averages of these six measurements were presented in the results section.

### Vascular cell adhesion molecule 1 (VCAM-1)

We used streptavidin-biotin method uses a biotin conjugated secondary antibody to link the primary antibody to a streptavidin-peroxidase complex for Immunohistochemistry (IHC) staining. The labeled streptavidin-biotin (LSAB) method were utilized to measure expresson of VCAM-1 in the aortic tissue of the rats. Firstly, aortic tissue were prepared and preserved through deparaffinize models following fixation. Secondly, aortic tissue were rehydrated by immersing the slides through the xylene (three washes 5 minutes each), 100% ethanol (two washes 10 minutes each), 95% ethanol (two washes 10 minutes each), 70% ethanol (two washes 10 minutes each), 50% ethanol (two washes 10 minutes each), and deionized water (two washes for 5 minutes). Thirdly, aortic tissue were washed using Phosphat Buffer Sollution and then, dipped into 3% of H_2_O_2_ solution withing 20 minutes. Fourthly, we added 1% of Bovine Serum Albumin to the Phosphat Buffer Sollution and then incubated them within 30 minutes in the room temperature. Fifthly, primary antibody anti-VCAM-1 (Santacruz biotech SC-13160) were added and incubated within 30 minutes, then washed again using Phosphat Buffer Sollution. Secondary antibody (Anti-Rat IgG Biotin Labelled) were added and incubated within 30 minutes in the room temperature, then washed using Phosphat Buffer Sollution. Sixthly, SA-HRP (Strepavidin-Hoseradish Peroxidase) complex were added and incubated within 10 minutes in the room temperature and then, washed using Phosphat Buffer Sollution. Seventhly, Chromogen DAB (3,3-diaminobenzidine tetrahydrochloride) were added and incubated within 10 minutes in the room temperature and then, washed using Phosphat Buffer Sollution and sterile water. Finally, counterstain Hematoxylin-Eosin were added into the object glasses and expression of VCAM-1 were measured and analyzed by a biological microscope (400x magnification) from tunica intima and tunica media of the aortic tissue. Semiquantitative measurements of VCAM-1 were done by immunoreactivity scoring system **(Table 1)**.

**Table 1:** Immunoreactivity Scoring System (IRS)

| Score for percentage of cells staining | Score for intensity of staining |
| --- | --- |
| 0 = no stained cells | 0 = no reaction |
| 1 = <10% cells are stained | 1 = mild intensity of staining |
| 2 = 10-50% cells are stained | 2 = moderate intensity of staining |
| 3 = 51-80% cells are stained | 3 = heavy intensity of staining |
| 4 = >80% cells are stained |  |

### Endothelial Nitric Oxide Synthase (e-NOS)

All samples were assessed by direct-sandwich enzyme-linked immunosorbent assay (ELISA) under the manufacturer’s (R&D System Europe Ltd, Abingdon, UK) according to the National Institute for Biological Standards and Controls (Blanche Lane, South Mimms, Potters Bars, Hertfordshire, UK) protocol. We used eNOS kit from the elabscience (catalogue number: E-EL-R0367). Briefly, samples from the aortic tissue were collected and stored at −70°C (−94°F) at Institute of Tropical Diseases Universitas Airlangga (UNAIR). Samples were homogenized into solution. Then, 100 μL of the solution were mixed with the well-coated primary antibody for e-NOS. Overnight incubation were done in the temperature 4°C with shaking machine. Wash Buffer (20x) were diluted to 1x working solution with D.I. water prior to ELISA wash procedures. After that, 50 μL of the stop solution were added into each samples. A minimum value of 0.01 pg/mL were assigned for below the limit of detection.

### Statistical analysis

All measurements were performed and replicated at least three times. Results were presented as (1) means ± standard deviations (SD) for normally distributed data; (2) medians with lower and upper value for abnormally distributed data. The assumption of the normality for the complete data was assessed by Shapiro-Wilk test. Test of homogeneity of variances was assessed by Levene Statistics. Statistical significance were examined by Independent T-test, Mann-Whitney U test, and logistic regression using SPSS version 17.0 for Microsoft (IBM corp, Chicago, USA).

## RESULTS

### Comparison of IMT level between smoke and non-smoke groups

After 28 days following experiments, there was a significance difference of IMT level between both groups (*p*<0.001). Mean of the aortic IMT in all subjects were 73.68±17.86 μm. Mean of the aortic IMT in cigarette smoke groups were 88.39±2.51 μm. Mean of the aortic IMT in control group were 58.98±13.61 μm. **Table 2** presents the impact of the exposure of daily 40 cigarette smokes on the aortic IMT profile of the experimental animals. The comparative analysis of IMT parameters demonstrated that there were a statistically significant differences between the groups (*p*<0.001; Mann-Whitney’s test). (**Figure 2**)

**Figure 2:**
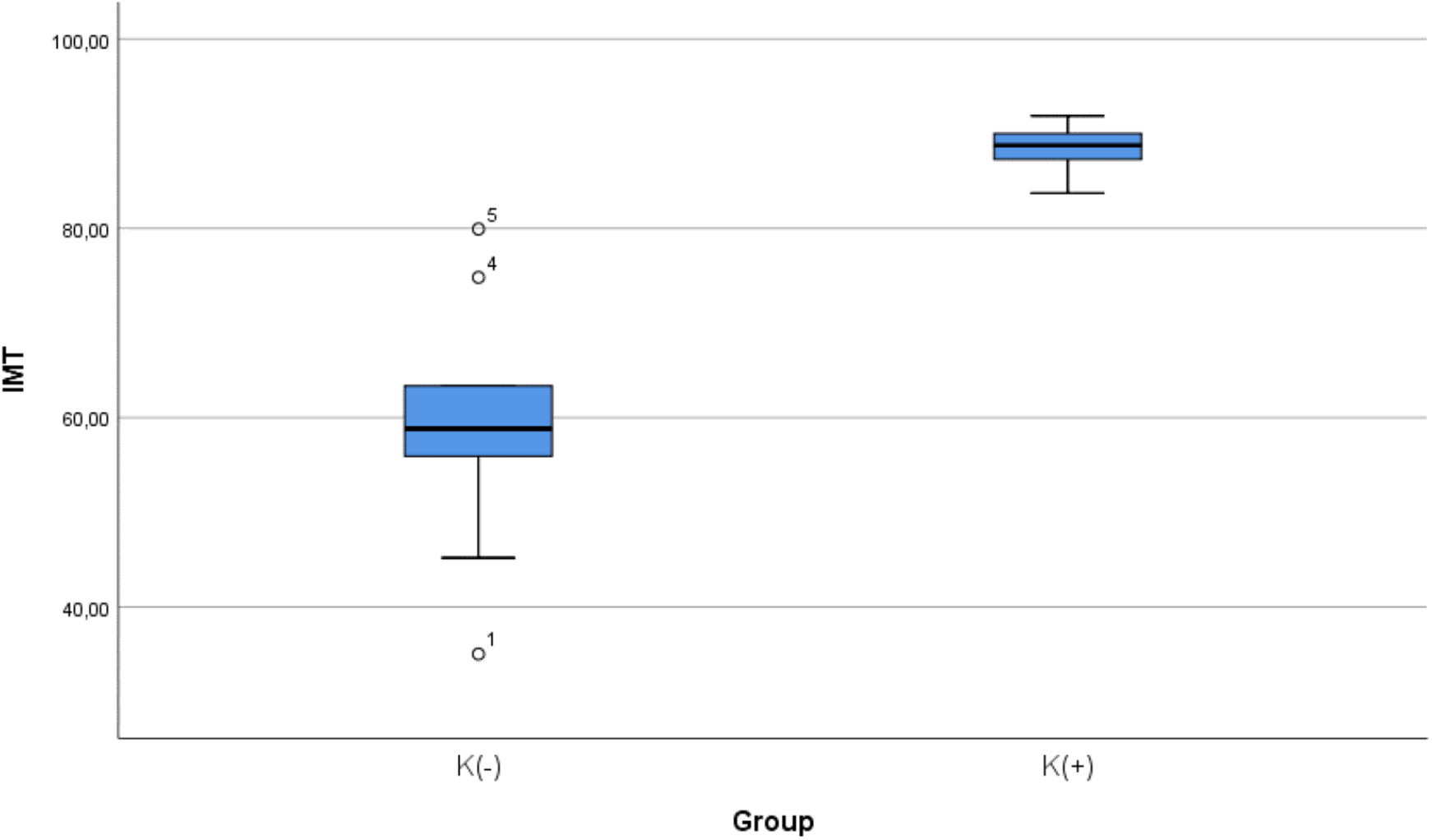
Median with lower and upper value of IMT between K(+) group which is exposed to the daily 40 cigarrete smokes and K(−) group as the control group.

**Table 2:** Statistic table IMT between K(+) group which is exposed to the daily 40 cigarrete smokes and K(−) group as the control group.

Group Statistics
|  | Group | N | Mean | Std. Deviation | Std. Error Mean | Sig (Independent T) | Sig (Mann-Whitney) |
| --- | --- | --- | --- | --- | --- | --- | --- |
| IMT | K(-) | 9 | 58,9800 | 13,61075 | 4,53692 | <0.001 | <0.001 |
|  | K(+) | 9 | 88,3911 | 2,51766 | ,83922 | <0.001 | <0.001 |

### Comparison of e-NOS level between smoke and non-smoke groups

After 28 days following experiments, there was a significance difference of e-NOS level between both groups (*p*<0.001). Mean of the e-NOS in all subjects were 78.02±25.84 pg/ml. Mean of the e-NOS level in cigarette smoke groups were 101.22±11.8 pg/ml. Mean of the e-NOS level in control group were 54.83±8.3 pg/ml. **Table 3** presents the impact of the exposure of daily 40 cigarette smokes on the e-NOS profile of the experimental animals. The comparative analysis of e-NOS parameters demonstrated that there were a statistically significant differences between the groups (*p*<0.001; Mann-Whitney’s test). (**Figure 3**)

**Figure 3:**
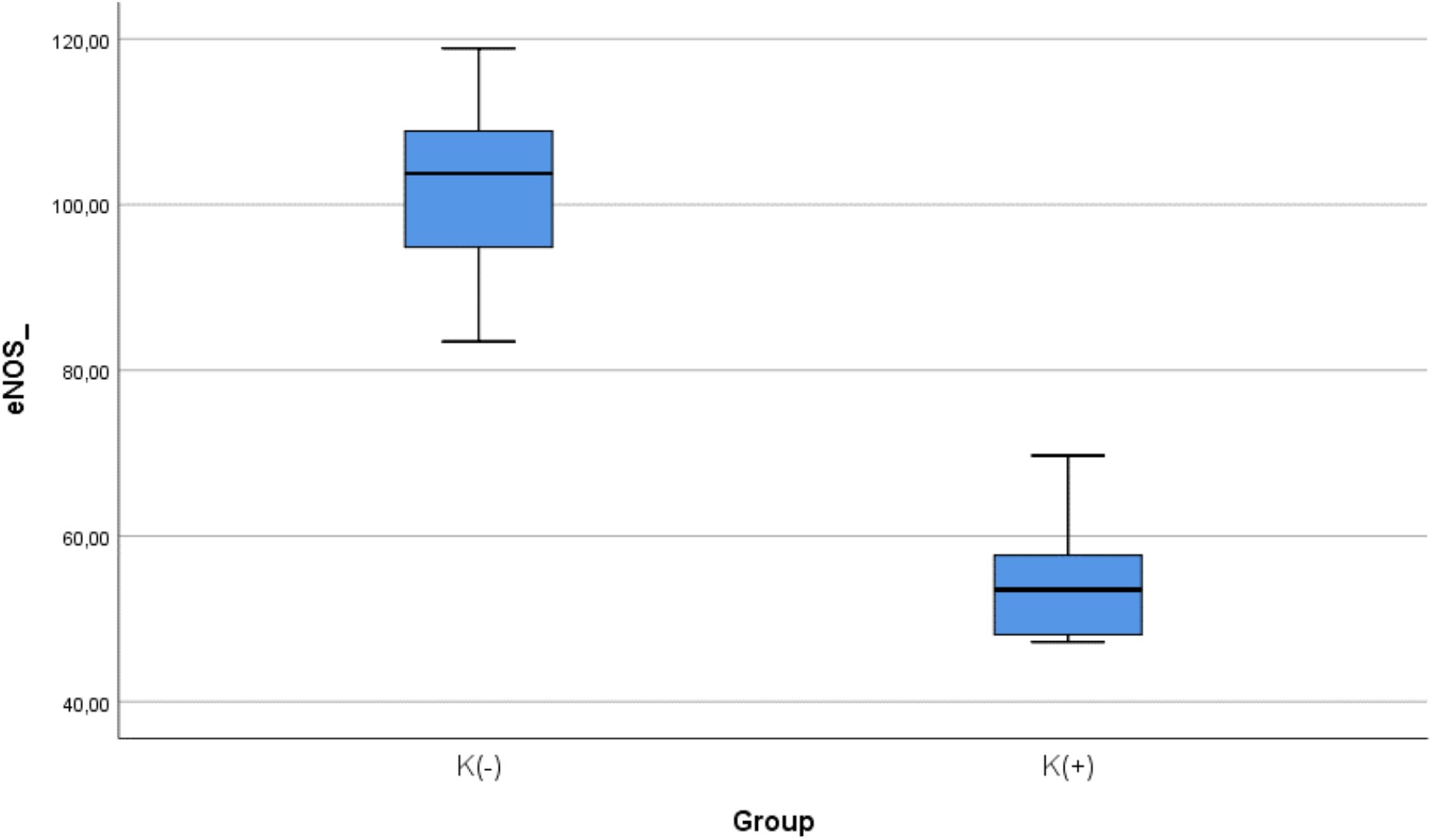
Median with lower and upper value of e-NOS between K(+) group which is exposed to the cigarrete smokes and K(−) group as the control group.

**Table 3:**
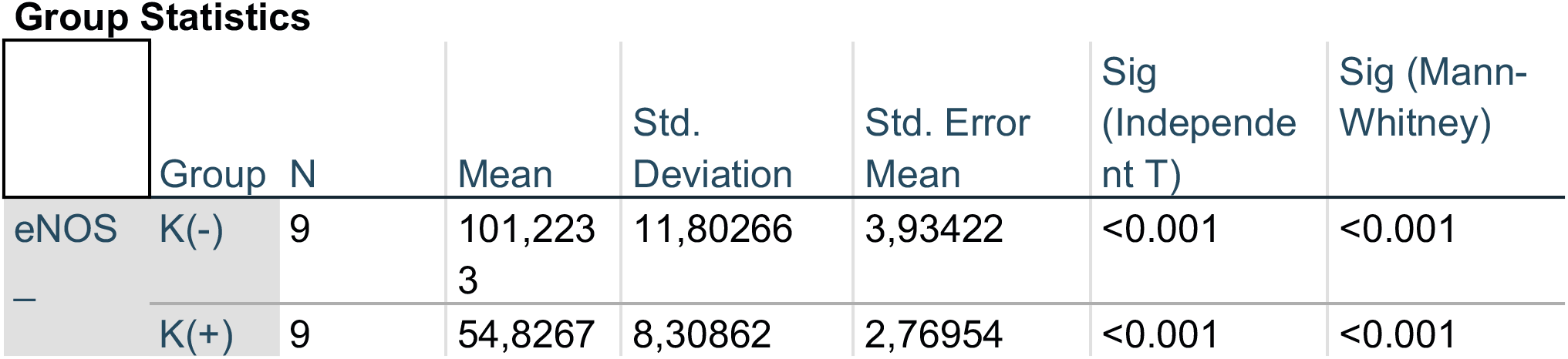
Statistic table e-NOS between K(+) group which is exposed to the cigarrete smokes and K(−) group as the control group.

### Comparison of VCAM-1 expression between smoke and non-smoke

After 28 days following experiments, mean of the VCAM-1 expression in all subjects were 9.00±3.51. Mean of the VCAM-1 level in cigarette smoke groups were 10.33±2.9. Mean of the VCAM-1 level in control group were 7.67±3.7. **Table 4** presents the impact of the exposure of daily 40 cigarette smokes on the VCAM-1 expression of the experimental animals. The comparative analysis of VCAM-1 expression demonstrated that there were no statistically significant differences between the groups (*p*=0.112; independent t test). (**Figure 4**)

**Figure 4:**
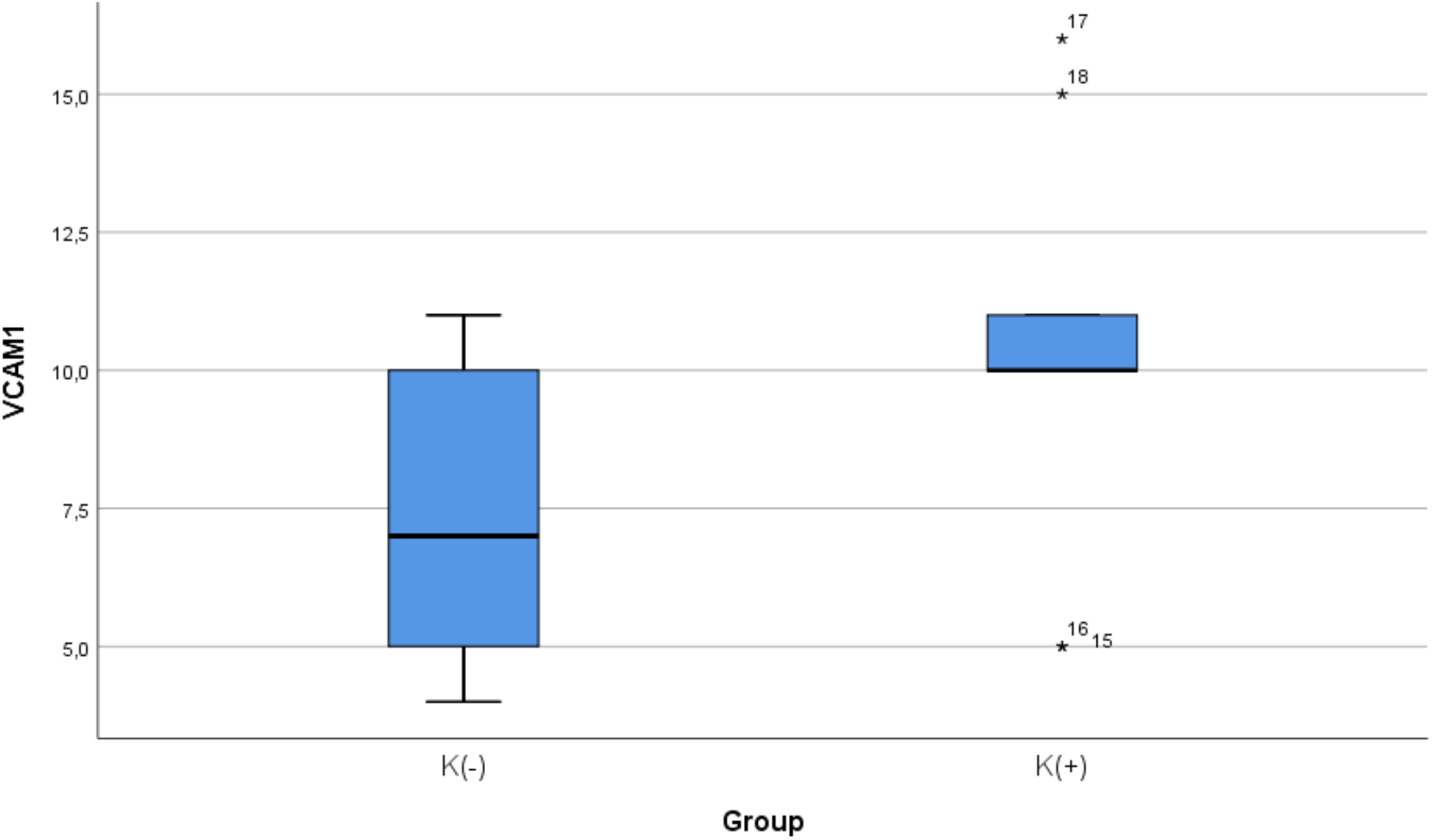
Median with lower and upper value of VCAM-1 between K(+) group which is exposed to the cigarrete smokes and K(−) group as the control group.

**Table 4:**
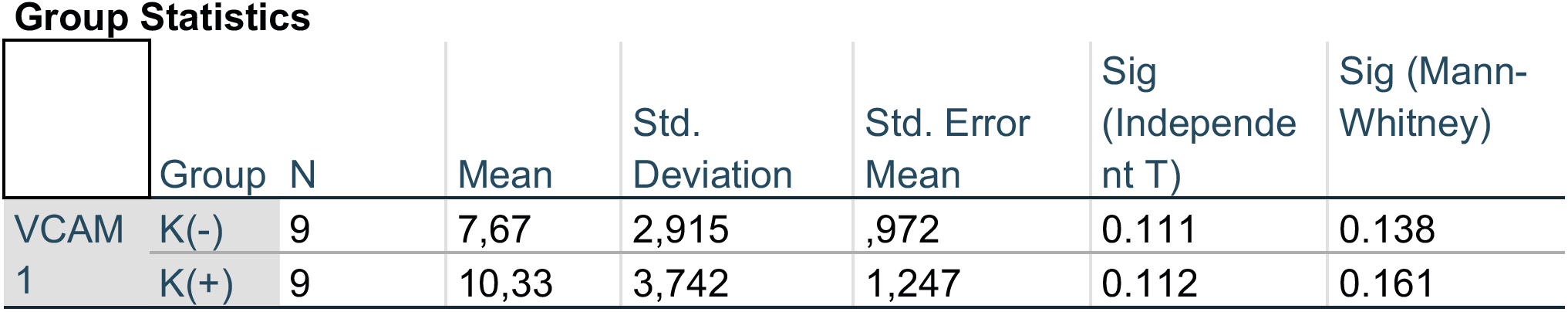
Statistic table of VCAM-1 between K(+) group which is exposed to the cigarrete smokes and K(−) group as the control group.

### Correlation of e-NOS level and aortic IMT

To determine if level of e-NOS is correlated with atherosclerosis, we measured e-NOS as a parameter of endothelial cell function in aortic tissue of Wistar rats. Linear regression model found that e-NOS was negatively correlate with aortic IMT in our experimental study (r^2^ = 0.584, β = −0.764, *p* < 0.001). (**Figure 5**)

**Figure 5:**
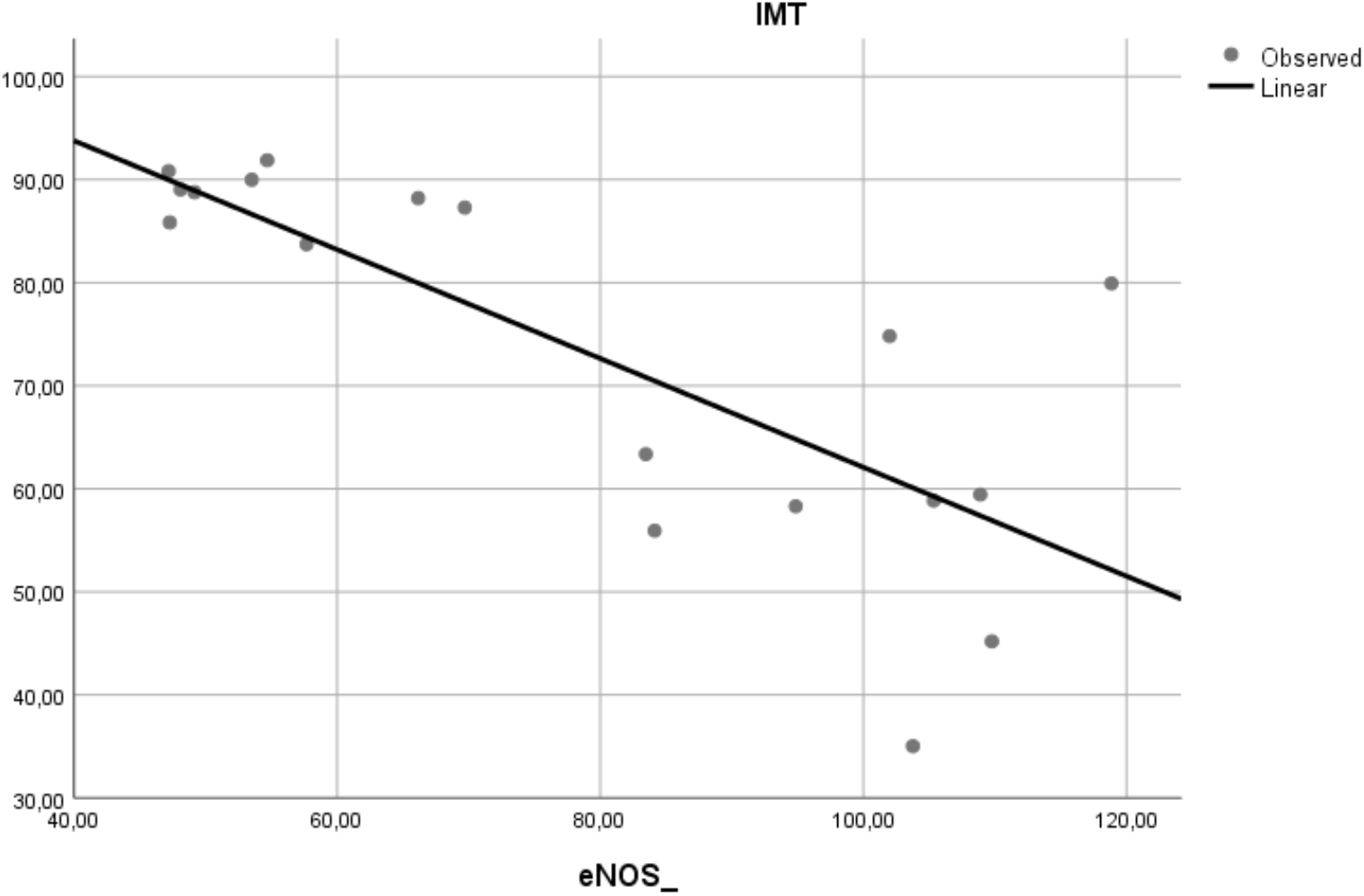
Relation between e-NOS level and aortic IMT in experimental rats. A negative linear relationship was found between e-NOS level and aortic IMT.

### Correlation of VCAM-1 expression and aortic IMT

To determine if expression of VCAM-1 precedes atherosclerosis, we measured expression of this adhesion molecule in aortic tissue of Wistar rats. Linear regression model found that VCAM-1 expression did not correlate with aortic IMT (r^2^ = 0.197, *p* = 0.065). (**Figure 6**)

**Figure 6:**
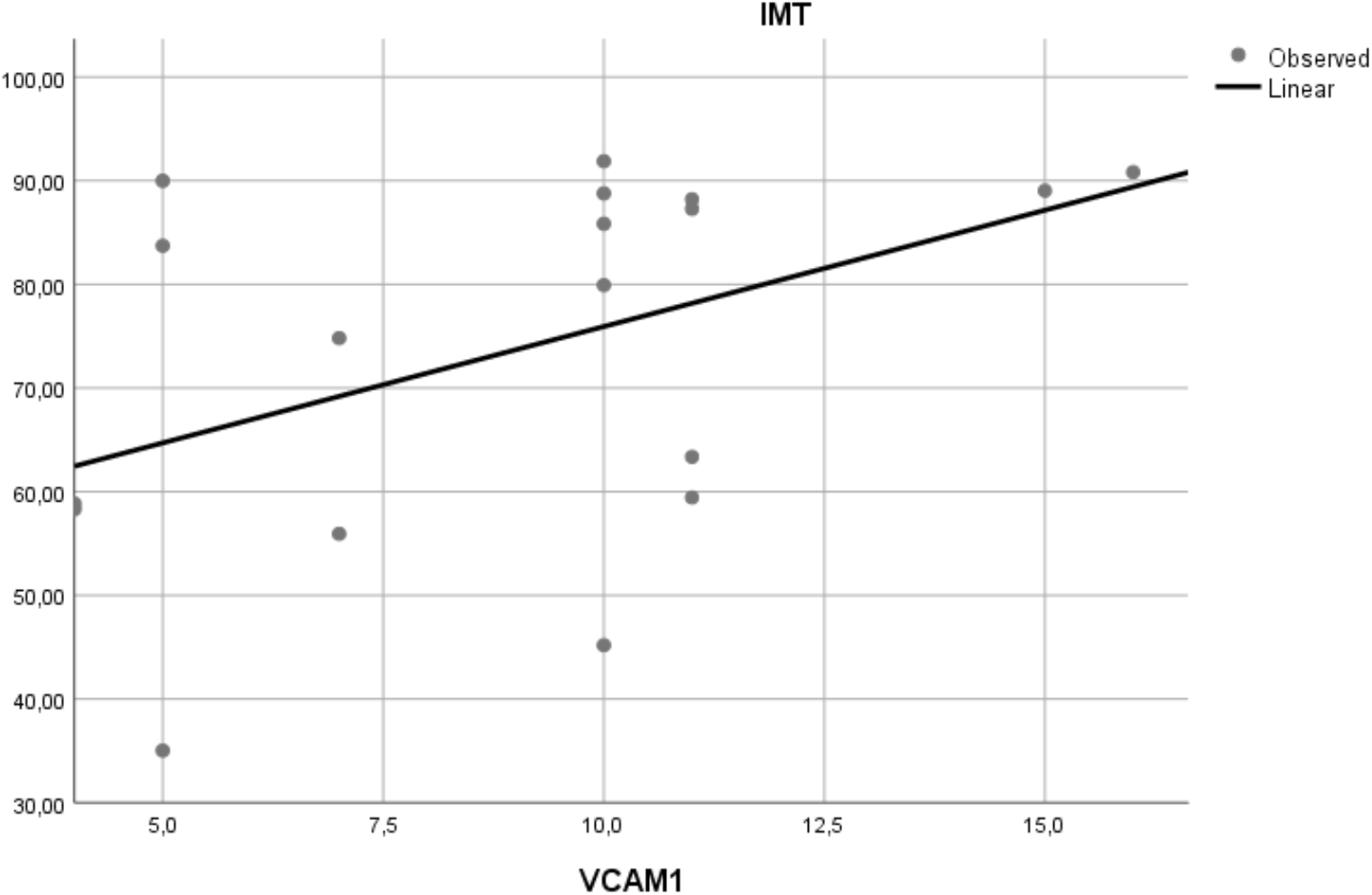
Relation between aortic VCAM-1 expression and aortic IMT in experimental rats. A positive but non-significant linear relationship was found between aortic VCAM-1 expression and aortic IMT.

## DISCUSSION

### Oxidative stress-mediated cigarette smokes precedes atherosclerosis

Cigarette smoking is one of the well-established modifiable risk factor for developing atherosclerosis, which mechanisms remain closely linked to the increased oxidative stress. Total amount of cigarettes smoked per day plays an essential role in increasing the level of oxidative stress and depletion of the antioxidant system. Cigarette smoke contains great concentrations of reactive oxygen species and tiny particles that are easily inhaled in human body^18^. It is believed that smoking causes increased oxidative stress because of several mechanisms, including direct damage by radical species and the inflammatory response caused by cigarette smoking. The production of oxidative stress and reactive oxygene species due to the cigarette smoke is expected to increase VCAM-1 expression and decrease of e-NOS level. According to the previous research by Yang et al (2014), an increase of VCAM-1 expression in rat arteries after being exposed to cigarette smoke had been observed for 7 days^19^. In a translational research did by Teasdale et al (2014) and Pott et al (2017) also supported that increased oxidative stress, reactive oxygene species, and VCAM-1 expression in endothelial cell cultures following exposed to cigarette smokes^20^. Previously, researchers had been studying the influence of smoking on the levels of several biomarkers of oxidative stress, antioxidant status and redox status, including plasma hydroperoxides, e-NOS and VCAM-1. Using different assays to our study, they confirmed that smokers have elevated concentrations of VCAM-1 and compromised e-NOS status^21^.

### Cigarette smoke extract induces expression of cell adhesion molecules

VCAM-1 is expressed in vascular endothelial cells, and expression of VCAM-1 may promote the adhesion of leukocytes to the endothelial cells. VCAM-1 accelerates the migration of adherent leukocytes along the endothelial surface, and promotes the proliferation of vascular smooth muscle cells; thus, VCAM-1 may play an essential role as a pro-atherogenic molecules^22^. Exposure to cigarette smoke in this study can increase VCAM-1 expression in the aorta although the increase is not statistically significant between the two groups. An insignificant increase in VCAM-1 expression was also found in the previous human research held by Noguchi (1999). In his previous research, soluble VCAM-1 levels were increased in smokers’ serum but not significantly when compared to non-smokers’ serum^23^.

Increased of VCAM-1 expression is a multifactorial process, smoking could not increase VCAM-1 independently without other risk factos such as dyslipidemia. Mu *et al* (2015) had proven this hypothesis by examining VCAM-1 expression in aortic tissue of dyslipidemia patients. As a result, VCAM-1 expression was positively correlated with triglyceride, total cholesterol and LDL levels while VCAM-1 and HDL had a negative correlation^24^. Because the expression of VCAM-1 in endothelial cells requires a trigger that is high lipid levels, especially LDL. An increase in oxidized LDL in the endothelium will be phagocytosed by macrophages. Recruitment of these macrophages requires the role of VCAM-1^25^. In our study, other factors contributing to the development of atherosclerosis such as dyslipidemia weren’t included. Our study did not use experimental animals with high-fat diets and serial lipid profile measurement Therefore, results of our study didn’t show any statistical significance of VCAM-1 expression between K (−) and K (+) groups.

### Cigarette smoke extract counteracts atheroprotective effects of endothelial nitric oxide synthase

Decreased bioavailability of NO is a central mechanism in the pathophysiology of endothelial dysfunction. Endhotelial nitric oxide synthetase (e-NOS) is an enzyme that resposible to produce NO in endothelial cells, so the level of eNOS can represent the availability of NO in endothelial cells^26^. Endothelial-cell dysfunction itself could be tested by acetylcholine response function and adenosine coronary flow reserve tests^27^. Celermajer *et al* (1992) published a study showing that smoking reduces flow-mediated dilatation (FMD) in systemic arteries in healthy young adults^28^.

Our study showed that exposure to cigarette smoke can reduce levels of eNOS in the aorta. Our results are consistent with the findings of Su et al and He et al. which shows a significant decrease of eNOS level in endothelial cell cultures exposed to cigarette smoke. He *et al* (2017) showed that exposure to cigarette smoke in endothelial cell culture can reduce the expression of eNOS genes and proteins, resulting endothelial-cell dysfunction^29^. On the other hand, Su *et al* (1998) had already proven that administration of cigarette smoke extract can reduce the expression of genes and proteins eNOS. The effect of eNOS reduction depends on the duration of exposure to the cells. The longer duration of cigarette smoke exposure, eNOS levels will be decreased^30^. In addition to decreasing eNOS at the gene level, Pini *et al* (2016) showed that exposure to secondhand smoke had also been shown to reduce eNOS at protein levels. eNOS levels decreased in the aorta of guinea pigs after exposed to cigarettes for 8 weeks^31^.

It has been demonstrated that cigarette smoking triggers demethylation, leading to a consecutive reactivation of epigenetically silenced genes in vitro and in vivo of eNOS and NO production^32^. Peroxinitrites, a very reactive oxygene species and pro-oxidant properties from cigarette extract, is believed to promote demethylation and inactivation of e-NOS^33^. In addition, peroxynitrite and other free radicals can deactivate BH4 which is an important cofactor in eNOS production. This was explained by the research of Abdelghany et al (2018) which showed that exposure to cigarette smoke has been shown to reduce the BH4 cofactor and correlated with the amount of superoxide and NO production in endothelial cell cultures^34^. A decrease of e-NOS and NO level will increase vascular tone, increase expression of adhesion molecules, and trigger coagulation cascade and inflammation^35^.

### In the final pathway, cigarette smoking leads to increase of aortic intima-medial thickness as an earlier sign of atherosclerosis

Based on these literatures and our own data, we suggest that the exposure to cigarette smoking for 28 days daily might be an independent risk factor for atherogenic process through several mechanisms. Aortic IMT in this study increased in group K (+) as was also found in studies conducted by Ali *et al* (2012)^36^. Increased aortic and entire blood vessels’ IMT are due to the pathological conditions such as apoptosis and excessive proliferation as a compensation mechanism^37^. In the previous study, increased of IMT is the complication of endothelial dysfunction leads to the atherosclerosis process^38^. Cigarette smoking exposure underlies the endothelial dysfunction by reduction of e-NOS level and increased of VCAM-1 expression^39^.

Exposure to cigarette smoke also affects the histological structure of the aorta. In this study, we found not only an increased of IMT, but also structural changes marked by disorganization and vacuolization of smooth muscle cells in tunica media of the aortic tissue. On the contrary, no changes were observed at the tunica intima. Exposure to cigarette smoke for 28 days in the study of Ali *et al* (2012) also found the same results: no changes at the tunica intima were observed from the experimental rat^36^. Another experimental study from Jaldin *et al* (2013) found that exposed to cigarette smoke for 8 weeks, only made a disorganization in vascular smooth muscle cells in tunica media^40^. Vacuolization is one of the complications from cytotoxic processes in the cells and earlier marker of preclinical atherosclerosis. Chemical components from the cigarette smokes can cause oxidative stress which is characterized by permanent vacuolization in cells. In the microscopic phenotyping, vacuolization makes vascular smooth muscle cells have different shapes and sizes, thus making cells become disorganized and lead to atherosclerosis^41^.

### Limitations and Strength

Every study has its limitations which emerge during the realization of the study, creates challenges and thus, should be highlighted. First, this study had limitations with regard to small number of samples which can increase the likelihood of error and imprecision. Second, results from animal model often do not translate into replications in human model. Level of e-NOS and VCAM-1 expression in Wistar rats are typically transient, whereas in human persists for many years. Other crucial difference is IMT, which is usually much lower in the Wistar rats than human. These factors may have an impact on the interpretation of our results. Thus, the findings should be interpreted within the context of this study and its limitations. The strengths of the study were its high statistical power and the homogeneity of each group to enable comparison between groups and periods.

### Conclusion

The present study indicates that, cigarette smoking adversely affects endothelial function and increases risk of atherosclerosis. Cigarette smoking as a risk factor for atherosclerosis is closely linked to the increased inflammatory process on the vascular endothelium. Low e-NOS level and high VCAM-1 level observed following smoke exposure may increase aortic IMT. Furthermore, smoking has also been found to influence the aortic IMT. Aortic IMT itself reflects the level of established CVD risk factors in apparently healthy men and women, adding to the evidence that cigarette smoking contributes to CVD through their inflammatory effects on the vascular endothelium.

## ACKNOWLEDGMENTS

The authors would like to acknowledge Hari Basuki Notobroto from Public Health Department Universitas Airlangga whose statistical expertise was invaluable during the analysis. The authors, hereby, acknowledge the above authorities and all staff, fellows, residents and laboratory assisstants from Department of Cardiology and Vascular Medicine, Faculty of Medicine Universitas Airlangga, who are willing to help in the technical aspect.

## CONFLICT OF INTEREST

All authors confirm that there are no conflicts of interest.

## AUTHOR CONTRIBUTIONS

Conceptualization: Ardiana M.

Project administration and funding acquisition: Ardiana M, Hermawan HO.

Data curation and formal analysis: Nugraha RA.

Investigation: Ardiana M, Hermawan HO.

Methodology: Ardiana M, Hermawan HO.

Resources and Software: Pikir BS, Santoso A.

Supervision and validation: Pikir BS, Santoso A.

Writing - original draft: Nugraha RA.

Writing - review & editing: Pikir BS, Santoso A

## AVAILABILITY OF DATA AND MATERIALS

The data that support the findings of this study are available from the corresponding author, upon reasonable request.

## CONSENT FOR PUBLICATIONS

Not applicable (public data).

## ABBREVIATIONS

ANOVA: analysis of variant
ELISA: enzyme-linked immunosorbent assay
e-NOS: endothelial Nitric Oxide Synthase
H_2_O_2_: Hydrogen peroxide
IACUC: Institutional Animal Care and Use Committee
IHC: Immunohistochemistry
IMT: Intima–media thickness
IRS: Immunoreactivity Scoring System
LSAB: Labeled Streptavidin Avidin Biotin
NIH: National Institutes of Health
PCR: Polymerase Chain Reaction
SA-HRP: Strepavidin-Hoseradish Peroxidase
SD: standard deviation
SEM: standard error of the mean
SPSS: Statistical Package for the Social Sciences
VCAM-1: Vascular Cell Adhesion Molecule-1

## Notes

http://repository.unair.ac.id/id/eprint/102952

### Competing Interest Statement

The authors have declared no competing interest.

